# Genetic variability and potential effects on clinical trial outcomes: perspectives in Parkinson’s disease

**DOI:** 10.1101/427385

**Authors:** Hampton Leonard, Cornelis Blauwendraat, Lynne Krohn, Faraz Faghri, Hirotaka Iwaki, Glen Furgeson, Aaron G. Day-Williams, David J. Stone, International Parkinson’s Disease Genomics Consortium (IPDGC), Andrew B. Singleton, Mike A. Nalls, Ziv Gan-Or

**Author notes:** denotes equal contribution (Mike A. Nalls, Ziv Gan-Or). denotes corresponding author.

## Abstract

**Background:** Improper randomization in clinical trials can result in the failure of the trial to meet its primary end-point. The last ∼10 years have revealed that common and rare genetic variants are an important disease factor and sometimes account for a substantial portion of disease risk variance. However, the burden of common genetic risk variants is not often considered in the randomization of clinical trials and can therefore lead to additional unwanted variance between trial arms. We simulated clinical trials to estimate false negative and false positive rates and investigated differences in single variants and mean genetic risk scores (GRS) between trial arms to investigate the potential effect of genetic variance on clinical trial outcomes at different sample sizes.

**Methods:** Single variant and genetic risk score analyses were conducted in a clinical trial simulation environment using data from 5851 Parkinson’s Disease patients as well as two simulated virtual cohorts based on public data. The virtual cohorts included a *GBA* variant cohort and a two variant interaction cohort. Data was resampled at different sizes (n = 200-5000 for the Parkinson’s Disease cohort) and (n = 50-800 and n = 50-2000 for virtual cohorts) for 1000 iterations and randomly assigned to the two arms of a trial. False negative and false positive rates were estimated using simulated clinical trials, and percent difference in genetic risk score and allele frequency was calculated to quantify disparity between arms.

**Findings:** Significant genetic differences between the two arms of a trial are found at all sample sizes. Approximately 90% of the iterations had at least one statistically significant difference in individual risk SNPs between each trial arm. Approximately 10% of iterations had a statistically significant difference between trial arms in polygenic risk score mean or variance. For significant iterations at sample size 200, the average percent difference for mean GRS between trial arms was 130.87%, decreasing to 29.87% as sample size reached 5000. In the *GBA* only simulations we see an average 18.86% difference in GRS scores between trial arms at n = 50, decreasing to 3.09% as sample size reaches 2000. Balancing patients by genotype reduced mean percent difference in GRS between arms to 36.71% for the main cohort and 2.00% for the *GBA* cohort at n = 200. When adding a drug effect to the simulations, we found that unbalanced genetics with an effect on the chosen measurable clinical outcome can result in high false negative rates among trials, especially at small sample sizes. At a sample size of n = 50 and a targeted drug effect of −0.5 points in UPDRS per year, we discovered 33.9% of trials resulted in false negatives.

**Interpretations:** Our data support the hypothesis that within genetically unmatched clinical trials, particularly those below 1000 participants, heterogeneity could confound true therapeutic effects as expected. This is particularly important in the changing environment of drug approvals. Clinical trials should undergo pre-trial genetic adjustment or, at the minimum, post-trial adjustment and analysis for failed trials. Clinical trial arms should be balanced on genetic risk variants, as well as cumulative variant distributions represented by GRS, in order to ensure the maximum reduction in trial arm disparities. The reduction in variance after balancing allows smaller sample sizes to be utilized without risking the large disparities between trial arms witnessed in typical randomized trials. As the cost of genotyping will likely be far less than greatly increasing sample size, genetically balancing trial arms can lead to more cost-effective clinical trials as well as better outcomes.

## Introduction

In the past few decades, clinical trials for neurodegenerative disease-modifying drugs have repeatedly failed. Between the years 2002 and 2012, 413 Alzheimer’s Disease (AD) trials were performed, with 99.6% resulting in failure.^1^ 83 of these trials were in phase III, which can cost an estimated $11.5 to $52.9 million.^2^ Success has also been elusive for Parkinson’s Disease (PD), where we find that drugs, such as Preladenant, that show potency in phase II often fail to be successful in phase III.^3^ Failures at this stage of clinical trials can be attributed to numerous reasons, but one reason for failure may be attributable to genetic risk variability and non-optimal randomization of patient trial arms that can create large sources of variation in genetic risk factors across trial arms. For example, high variance between patients across trial arms led to trial failure for two solanezumab trials, where post-hoc analysis revealed that out of the 27% of patients with confirmed amyloid biomarker status, 25% of mild AD patients and 10% of moderate AD patients were amyloid negative, and so did not show typical progression of AD.^4^ Only in post-trial analysis was solanezumab found to be most effective for mild AD patients with higher rates of disease progression and less effective for mild patients in the lowest progression percentiles (atypical AD).

As most clinical trials lack genetic balancing, the resulting disease heterogeneity due to underlying genetic risk factors creates variability between individual patients and between overall trial arms in areas such as disease progression, which may be contributing to the high failure rate. This disease heterogeneity can be seen when examining well-known Parkinson’s disease risk variants, such as those in the *GBA* and *LRRK2* regions, which can alter disease etiology depending on the variant present in that participant.^5,6,7^ Polygenic risk scores for Parkinson’s have also been connected to disparity in disease etiology, with an increase in risk score corresponding in a decrease in disease age at onset.^8^ Incorporating genetic data into clinical trials has been shown to be beneficial, as is seen in a meta-analysis of clinical data from failed trials investigating the efficacy of lithium carbonate on amyotrophic lateral sclerosis (ALS) patients. After genetic analysis, there was a potential positive result for patients with a specific risk genotype (*UNC13A*). Not only does this show that genetic variation can influence trial outcome, but the study also suggests that unbalanced risk allele variability may be influencing false positive and false negative rates. They note a 24% chance of a genotype imbalance greater than 10% if the genotype in question is present in 15% of the cases at a sample size of n = 50.^9^ This idea leads us to the purpose of this paper, to explore in detail the disparity between trial arms due to variability in overall cumulative genetic risk and single variants, necessitating the collection of genetic data to balance samples pre-trial, or at the minimum, post-trial analysis in order to find previously hidden effects.

For Parkinson’s Disease, motor or cognitive symptoms serve as measurable outcomes in clinical trials. One commonly used measure, the Unified Parkinson’s Disease Rating Scale (UPDRS), assesses the severity of PD symptoms as well as functioning as a readout to measure drug efficacy. However, genetic heterogeneity can cause disparity in terms of the progression and presentation of PD symptoms, potentially affecting overall UPDRS readings, and thus, clinical trial outcomes. A study investigating predictors in motor progression in PD patients found that an interaction effect between two SNPs, rs9298897 and rs17710829, resulted in a ∼2 point increase in MDS-UPDRS score per year, indicative of a faster rate of motor decline in those patients.^10^ Other studies have also found associations between certain variants and their effect on PD symptoms. For example, different mutations within the *GBA* gene lead to differential effects on PD phenotypes.^5,6^ Carriers of severe *GBA* mutations have an age of onset (AAO) roughly 5 years earlier and around a 3-4 fold increase in disease risk, compared to mild *GBA* mutation carriers.^11^ Another example of this is seen among *LRRK2* mutations, with different variant possessing different molecular mechanisms and cellular effects. *LRRK2* G2019S, for example, is involved with kinase activation and lysosomal positioning alteration, where *LRRK2* R1441C is linked to GTP hydrolysis disruption and has no known effects on lysosomal positioning.^7^

To say that all Parkinson’s Disease clinical trials do not undergo some extent of pre-trial genetic adjustment would be incorrect, as there are clinical trials underway specifically for PD patients who carry a *GBA* or a *LRRK2* mutation.^12,13^ However, even within these specific subgroups of PD mutations, large variation between patients still exists. *GBA* is a prime example, where as discussed above, different *GBA* mutations can differentially affect both the risk and AAO of PD.^11,14^ The varying effect estimates of these different variants can also be used to quantify an individual’s disease risk into a total polygenic or genetic risk score (GRS). A study investigating the relationship between AAO and GRS comprised of dozens of GWAS risk variants found that a single standard deviation increase in patient GRS resulted in a 37 day decrease in AAO.^8^ Other studies suggest that a single SD increase in nearly the same GRS may speed onset to almost 1 year earlier.^15^ The difference in these findings can likely be attributed to heterogeneity in data across dozens of sites, as measurement and recording of AAO can be imprecise. However, the relationship between AAO and PD symptoms are well described in many studies, finding that variance in AAO leads to variance in mortality as well as variations in presentation of motor and non-motor phenotypes.^16,17^ If trial arms are significantly different in terms of GRS and single risk variant distribution, this will affect how the phenotypes of each arm and individual will present over time, which may affect the overall outcome of a trial. As a variety of variants affect disease characteristics like AAO or important markers of progression such as UPDRS, balancing trials on a genotype level will create better-matched trial arms. Genetic risk factors can also be useful as selection criteria for enrollment in a trial, with some studies already suggesting that certain genotypes, such as the ApoE genotype, be used for stratification or as covariates.^18^ Using the genetic information of patients to balance or build a trial can reduce heterogeneity that may result in better trial outcomes.

In this study, we used data from a large PD cohort to investigate genetic risk variability as a result of classical randomized trial simulations. The type of variance we focused on is related to two different genetic effects. We also investigated variance in virtual cohort simulations, using an interaction effect of two variants on a measurable clinical outcome to determine false negative and false positive rates. An additional virtual cohort was created to determine if patterns within a small subgroup of large-effect *GBA* risk variant carriers reflect that of the larger cohort with a wider range of mutation or variant types and locations. As variation in genetic load and variant distribution creates differences in disease phenotype, these simulations attempt to quantify variability in manifest disease symptoms that could serve as confounding variables in a clinical trial read out, possibly affecting the chance to determine a true treatment effect.

## Methods

### Genetic risk score and variant nomination from GWAS

A genetic risk score for each patient was calculated from the cumulative effect of each of the 47 variants nominated by GWAS.^19^ The regression coefficient for each allele determined the contribution to each patient’s overall risk score in terms of the allele dose of that variant, which was converted to possible counts of 0, 1, or 2. The formula as used and explained in Chang et al, 2017^19^, is below:

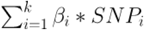

In the above formula, k represents the total number of variants, β _i_ is the regression coefficient associated with the effect allele from the GWAS, and SNP _i_ is the variant. This formula is applied to all PD patients and controls in the dataframe, which results in a cumulative risk score calculation for each person. GRS of all observations are then scaled to Z-scores (standard deviations of risk) weighted by the controls. Mean imputation was used for missing variants.

### Single Variant and Genetic Risk Score Analysis

Data from 5851 PD patients was sampled for 1000 iterations at sizes of 200, 400, 600, 1000, 2000, 3000, 4000, and 5000 observations. This cohort can be obtained from dbGaP^20^ and is well described in Nalls et al, 2014.^21^ Individuals were assigned to simulated treatment and placebo arms in a 1:1 ratio by a simple procedure of alternating individuals to each arm as they are sampled from the larger population. Binomial generalized linear models employing logit link functions were then used to classify whether individual patients belonged to either the randomized treatment or placebo arm using each of the 47 nominated variants as predictors. Variant significance level for association was calculated at every iteration of each sample size for each of the 47 variants individually, in combination with age and sex as well as 5 principal components as covariates. As our goal was to study the effects of random differences between trial arms, significant variants were determined by a significance threshold of p < 0.05. Magnitude of allele distribution difference was calculated by finding the allele dosage for each of the 47 variants per each trial arm, and then computing the percent difference in allele carrier status for each variant for 1000 iterations. These values were then averaged to represent the mean percent difference across all 47 variants for each sample size, both for significant iterations and all iterations. Instances where one trial arm had patients that possessed a certain variant and the other trial arm had no patients with that variant were removed, as percent difference calculations are not meaningful in this case, making our models slightly more conservative concerning rarer variants in small sample sizes.

In addition to analysis of cumulative allele frequencies at specific SNPs of interest, we also investigated difference in mean GRS between trial arms. The GRS analyses are quite similar to those looking at cumulative allele frequencies, however the GRS analyses take into account weights from previously published risk estimates. To determine significant differences in mean GRS between trial arms, a Z-score was calculated using a two sample Z-test at every iteration. The distribution of the GRS was non-normal, however, the two sample Z-test allows non-normal distributions if large enough samples are used. All sample sizes were large enough to allow sample variance to be used in Z-score calculation. Iterations were then filtered by significance of their Z-score, either falling above or below the 95% significance cut-off values of ± 1.96. For both significant iterations and all iterations combined, percent difference between the mean GRS of the treatment and placebo arms was calculated and these values were then averaged to obtain an overall mean difference in GRS for each sample size. A basic visualization of the workflow for GRS and single variant analysis is provided for further clarity. (Figure 1).

**Figure 1.**
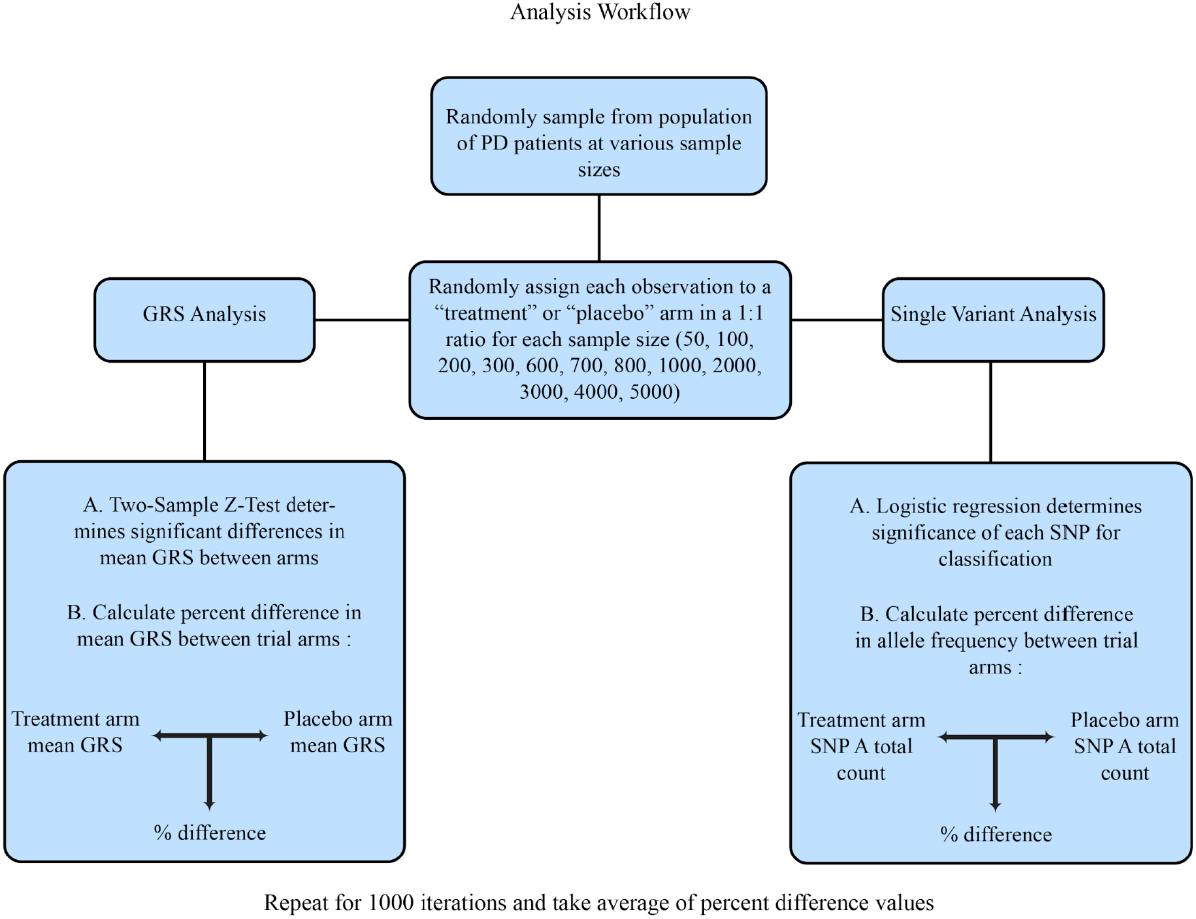
Analysis Workflow. A visualization of the analysis process. This workflow was followed for every chosen sample size and repeated for 1000 iterations.

Disparity in variance in GRS between arms was also investigated, as variance between two arms of randomly assigned patients will not always be equal. Trial arms were compared by performing a Levene’s Test for Equality of Variances at each iteration. Additional description of methods for this section of analysis focusing on within group variance estimates can be found in the supplemental material.

After initial variance analysis of completely randomized cohorts, an algorithm was used to balance patients between trial arms by genotype using the 47 variants nominated by GWAS. The algorithm was designed to imitate rolling admission to clinical trials by assessing the best placement (placebo or treatment arm) for each individual patient as they were added to the simulated trial. Best placement of each patient was determined by minimizing difference in overall allele dosage between trial arms.

### Virtual Cohort Simulation: rs9298897 and rs17710829

In a study by Latourelle et al, two variants (rs9298897 and rs17710829) had a replicated interaction effect of a roughly additional 1-2 point increase in MDS-UPDRS (parts II and III) per year.^10^ To create a virtual cohort of carriers, these variants were assigned to a population of 5000 according to Hardy-Weinberg equilibrium and European allele frequency estimates as reported by the ExAC Release 1 database.^22^ The estimates from this database are slightly lower than what would be seen in PD cases, and are therefore more conservative. Change in UPDRS score from baseline was used to determine significant difference between arms. As change in UPDRS was the chosen metric, the initial score for each patient would not affect simulation results, however a range of baseline scores were chosen in order to mimic conditions in real trials. All virtual patients were randomly assigned a baseline UPDRS score on a range from 15 to 25, such that scores within that range followed a uniform distribution. For each of the two simulated years, all virtual patients were assigned a random progression in MDS-UPDRS score of either 1 or 2 points per year, a more conservative progression rate based upon the average increase in MDS-UPDRS scores found in the Holden et al study^23^, for simulation purposes. Carriers of both the rs9298897 and rs17710829 variant received an additional increase in UPDRS score in accordance with the model effect size reported in Latourelle et al (β = 2.374, SE = 0.436). This cohort was sampled at sizes of 50, 100, 200, 300, 600, 700, and 800 observations and patients were randomly assigned to simulated treatment or placebo arm. Both false positive and false negative rates were investigated in this stage by performing 2 sample Z-tests for each iteration. False positive rates were determined by investigating the number of significant iterations that occured without any simulated “drug” effect. Percentage of false positives caused by the addition of the SNP interaction effect were calculated by comparing results of tests with and without the effect. For false negative rates, a simulated “drug” effect that decreased UPDRS score by 0.5 points per year was added to the patients in the treatment arm.

### Virtual Cohort Simulation: GBA

A virtual cohort for *GBA* mutation carriers was generated for simulation use. Many of the variants associated with this gene have low allele frequencies, resulting in only a small amount of real data. A virtual cohort was created to counteract this problem and create estimates for the behavior of variance within this cohort.

Three of the 47 variants used in this study are *GBA* variants, and these same variants were used to create a virtual cohort of patients, representing one of the many ongoing or upcoming *GBA* focused interventional trials. Using effect estimates from this study, individual genetic risk contribution was assigned to each variant. The variants were p.N370S (rs76763715) (β = 0.747, 95% CI [0.60, 0.90]), p.E326K (rs2230288) (β = 0.636, 95% CI [0.55, 0.72]), and p.T369M (rs75548401) (β = 0.362, 95% CI [0.23, 0.50]), all three of which have been associated with risk for PD.^24,25,26,27^ Each variant was then assigned to a population of ∼60,000 according to Hardy-Weinberg equilibrium and European allele frequency estimates as reported by the ExAC Release 1 database.^22^ This cohort was created in such a way that the individual GRS of each virtual patient was created by the combination of the three chosen *GBA* variants alone. Patients were filtered for those who possessed at least one of the chosen mutations and then sampled at sizes of 50, 100, 200, 300, 600, 700, 800, 1000, and 2000 observations to simulate *GBA* targeted trials. Raw GRS scores were used for analysis of this cohort rather than control-weighted Z-scores as with the larger cohort. Average percent difference in GRS between trial arms was calculated for each sample size.

### Polynomial Regression, Locally-Weighted Polynomial Regression, and PCA

Locally-weighted polynomial regression models were trained on the mean percent difference in GRS and variance in GRS across trial arms for each sample size. LOESS models rely on weighted least squares and a smoothing parameter to fit a polynomial to different subsets of data, allowing a nonparametric and flexible fit to non-linear relationships. The models estimated best fit lines for the relationship between percent difference in GRS variance and sample size, and confidence intervals were calculated for each fit. These models were used to provide insight into the trends in the data and model fit. Additional polynomial models (non-locally weighted) were also trained to garner further information about variable significance in the relationships. In addition to regression models, principal component analysis (PCA) was performed using the variant dosage status of all 47 variants at a sample size of 200 for balanced and unbalanced cohorts. Logistic models were made using the components of this analysis as predictors for determining classification of a patient to either the treatment or placebo arm. PCA was also used to generate a visual example of the effects of balancing by plotting component scores from the first 3 components for both balanced and unbalanced cohorts at a sample size of 200. All statistical and modeling analyses were conducted with R.^28^ Code is available to the public through the National Institute on Aging Laboratory of Neurogenetics Github at https://github.com/neurogenetics/Clinical-Trial-Outcomes.

### Variance Using Heritability Estimates

Current heritability estimates were used as proxies to adjust the variance levels found in the clinical trial simulations. As the collective knowledge of the genetic causes of PD is far from complete, we took PD genetic heritability estimates into account to provide variance estimates on two ends of the spectrum. On the top or “ceiling” end of the spectrum, we assumed a one-to-one relationship between phenotypic variance and the underlying genetic variance in the patients. Therefore, the values obtained from the clinical trial simulations would represent the expected phenotypic variance levels between arms in similar, real-world trials. On the low or “floor” end, we looked at variance in terms of the genetic information currently available to us. This “floor” version is the most realistic estimate at present, as it depends on our current knowledge of risk genetics related to PD. We can only balance genetics in trials to the extent of what we know is correlated with the disease in question, and thus the “floor” values are representative of the expected phenotypic variance between arms based upon the information known to us. This analysis was performed using difference in mean GRS results from the clinical trial simulations and the current genetic heritability estimate for PD, which is ∼26.9%.^29^ We then calculated the “floor” values by looking at the percent difference in variance at each sample size as a proportion of the heritability estimate.

## Results

### High Genetic Heterogeneity with Randomization of Different Simulated Trial Sizes

To examine how randomization of patients at different sample sizes affects variability in overall genetic risk score disparity between arms, we performed 1000 iterations of sampling and randomization of trial arms for different sample sizes. We then calculated the mean difference in GRS as a percentage, between the treatment and placebo arms in significantly different iterations and across all iterations regardless of significance. Evaluation of GRS differences across trial arms revealed that overall percent difference between trial arms was high at small small sample sizes, and the magnitude of difference decreased as sample size increased. (Figure 2.A.). Results from analysis of differences in variance in GRS between trial arms were similar to differences in mean GRS, with a high percent difference between arms that decreased as sample size increased. Additional results and tables for this can be found in the supplemental material. Iterations significant in either variance or mean GRS difference between arms accounted for roughly 10% of iterations at all sample sizes. Percent differences in GRS ranged from close to 0 to over 100%. Significantly different iterations showed higher rates of difference in GRS as expected between simulated treatment and placebo arms. At smaller sample sizes, percent difference in GRS score can be over 100% when comparing trial arms, with this difference decreasing to roughly half that amount as the sample size reaches 1000 patients. (Table 1).

**Figure 2.**
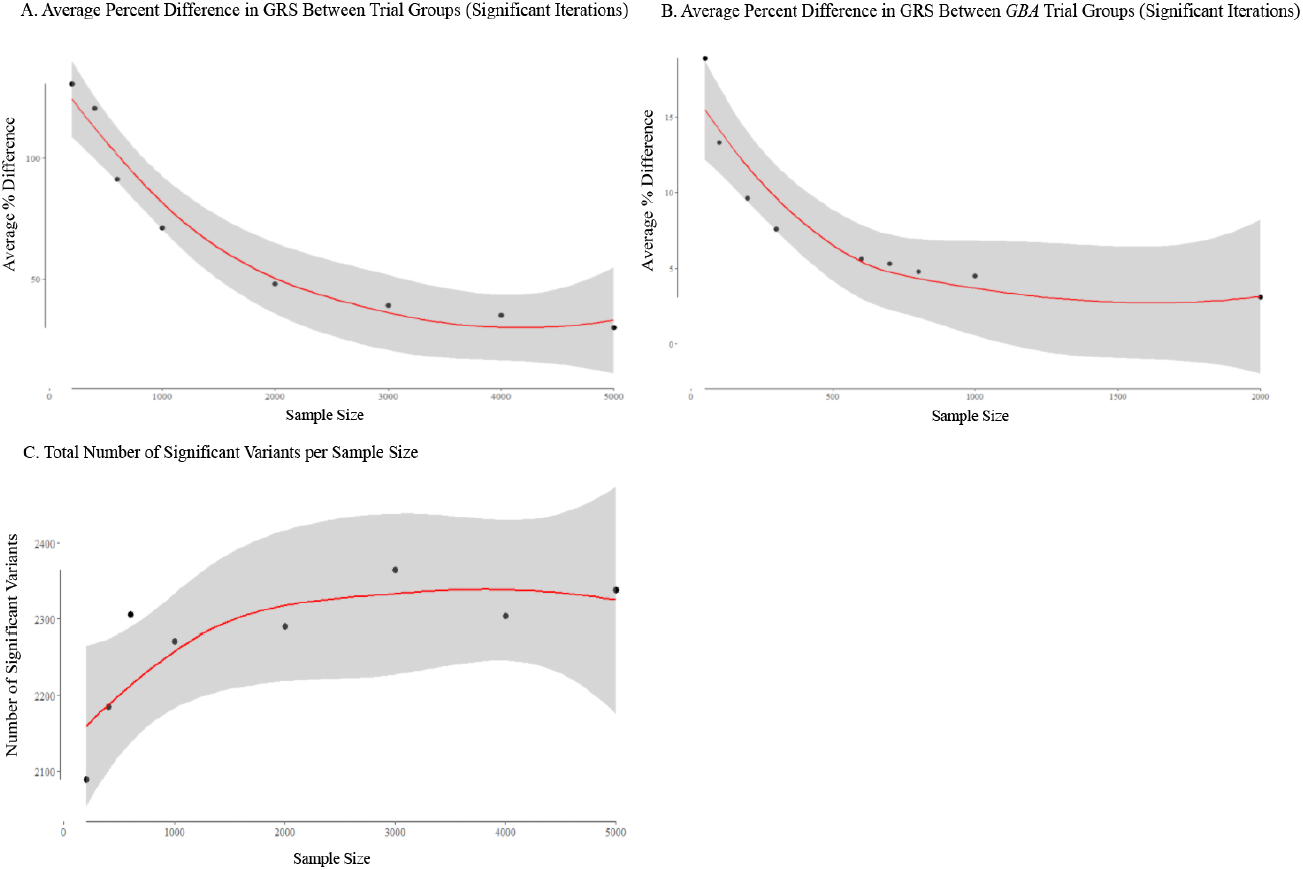
Average percent difference in GRS between and within trial groups; number of significant variants per sample size. Lines are fit to relationships using locally-weighted polynomial regression models (LOESS) and confidence intervals are shown in grey. Residual standard error and span values were obtained from the LOESS models and significance and R^2^ values were obtained from polynomial regression models. (A) The rehtionship between percent difference in between group variance and sample size is shown. Signifcant iterations were determined by Z score cut-off values of ± 1.96 (p <0.05). Span = 1.4; Residual Std. Error = 7.946; Adjusted R^2^ = 0.928; P-value = 0.0006. (B) The relationship between percent difference in between group variance and sample size is shown for the *GBA* cohort. Signifcant iterations were determined by Z score cut-off values of ± 1.96 (p < 0.05). Span = 1.17; Residual Std. Error = 2.379; Adjusted R^2^ = 0.794; P-value = 0.0032. (C) The relationship between the total number of times a variant was a significant predictor and sample size is shown. Significance for prediction was determined by a binomial generalized linear model with a logit link function (p <0.05). Span = 1.4; Residual Std. Error = 57.52; Adjusted R^2^ = 0.585; P-value = 0.097.

**Table 1.** Average % Difference is Mean GRs

| Sample Size | Average % Difference in GRS (CI 95%) Significant Iterations | Average % Difference in GRS (CI 95%) All Iterations | % Difference Range (Min - Max) |
| --- | --- | --- | --- |
| 200 | 130.87 (120.89 - 140.85) | 51.55 (49.01 - 54.10) | 0.079 - 198.76 |
| 400 | 108.47 (99.10 - 117.85) | 36.60 (34.77 - 38.42) | 0.045 - 179.82 |
| 600 | 91.31 (85.58 - 97.05) | 29.14 (27.67 - 30.61) | 0.024 - 146.82 |
| 1,000 | 71.15 (66.91 - 75.39) | 22.64 (21.52 - 23.77) | 0.006 - 103.84 |
| 2,000 | 48.03 (45.66 - 50.40) | 16.67 (15.86 - 17.48) | 0.006 - 86.71 |
| 3,000 | 39.00 (37.45 - 40.56) | 13.91 (13.25 - 14.56) | 0.001 - 56.55 |
| 4,000 | 35.02 (33.20 - 36.84) | 11.22 (10.69 - 11.75) | 0.016 - 53.26 |
| 5,000 | 29.87 (28.43 - 31.31) | 10.58 (10.09 - 11.07) | 0.014 - 47.12 |

The average percent difference between trial groups and related confidence intervals are shown for each sample size for both significant iterations and total iterations. Significant iterations were determined by Z score cut-off scores of $\pm 1.96$ ( $p < 0.05$ ). Minimum and maximum percent differences found between groups for all 1000 iterations are also shown.

### Variation in Single Variant Distribution Between Randomized Arms

Single variant analysis revealed that approximately 90% of the trials, regardless of trial size, resulted in a significant difference in allele frequency of the risk SNPs between treatment and placebo arms. We found that number of significant variants (unadjusted P < 0.05) fluctuated with sample size. This suggests that it is unlikely that simply increasing sample size will result in a reduction in the number of significantly differently distributed variants between arms. This is a function of allele frequency and statistical power as sample size increases. In addition, there was a non-significant correlation between sample size and number of significant variants (r = 0.686, p = 0.061), that suggests that increasing sample size may result in an overall increase in the number of significantly different variant distributions (Figure 2.C.). However, while number of significant variants may increase, the percent disparity in allele frequency of the SNPs of interest between arms decreases. For significant iterations, average percent difference in cumulative risk allele dosage decreases from 41.60% to 27.60%, a drop in difference of 14% (p = 5.76e-66, 95% CI [12.42, 15.57]) as sample size increases from 200 to 1000. (Table 2.) True percent difference between arms is likely higher than stated here, as situations where either one of the trial arms possessed zero counts of a rare variant could not be included in percent difference calculations.

**Table 2.** Average % Difference in Variant Dosage Between Groups

| Sample Size | Average % Difference (CI 95%)<br>Significant Iterations | Average % Difference (CI 95%) All<br>Iterations | % Difference<br>Range (Min - Max) |
| --- | --- | --- | --- |
| 200 | 41.60 (40.57 - 42.62) | 19.80 (15.50 - 24.11) | 0 - 176.471 |
| 400 | 33.80 (32.68 - 34.93) | 15.97 (11.52 - 20.41) | 0 - 166.667 |
| 600 | 30.40 (29.27 - 31.52) | 13.61 (9.51 - 17.71) | 0 - 175.000 |
| 1,000 | 27.60 (26.41 - 28.80) | 10.75 (7.36 - 14.14) | 0 - 166.667 |
| 2,000 | 21.28 (20.31 - 22.25) | 5.03 (3.45 - 6.62) | 0 - 129.41 |
| 3,000 | 17.29 (16.53 - 18.05) | 6.18 (4.25 - 8.11) | 0 - 131.03 |
| 4,000 | 15.26 (14.59 - 15.94) | 5.34 (3.66 - 7.03) | 0 - 97.44 |
| 5,000 | 13.81 (13.19 - 14.43) | 4.81 (3.28 - 6.35) | 0 - 92.86 |

The average percent difference in allele dosage for each of the chosen 48 variants between trial groups and related confidence intervals are shown for each sample size for both significant iterations and total iterations. Significant iterations were determined by FDR corrected p values ( $p < 0.05$ ). Minimum and maximum percent differences found between groups for all 1000 iterations are also shown.

### Hypothesized False Negative and False Positive Rates

A replicated interaction effect between two variants associated with an increase in MDS-UPDRS was used to demonstrate the effects of imbalanced trials on overall outcome. We found that at small sample sizes, 33.9% of trials resulted in a false negative with the simulated drug effect of a 0.5 point reduction in UPDRS score per year at n = 50. False negative rate decreased as sample size increased, reaching nearly 1.0% as sample size approached 200. With the addition of the second year of the trial, percentage of false negatives decreased across sample sizes, however, 21.2% of iterations still resulted in a false negative at n = 50. Number of iterations that resulted in false positives and false negatives were compared both with and without the interaction effect. This allowed us to determine how many false positives and negatives were truly attributable to in imbalance of these SNPs between arms. Percentage of false negatives caused by the SNP interaction effect alone increased with sample size, with 100% of the observed false negatives being caused by the interaction as sample size increased to 200 for trial year 1. Nearly 100% of false negatives in trial year 2 were caused by the SNP interaction effect alone. (Table 3). False positives and the percentage of false positives caused by the SNP interaction effect were fairly similar across sample sizes. (Table 4).

**Table 3.** False Negative Rates for Unbalanced Trials Over a Simulation of 2 Years

| Sample Size | Year 1 |  | Year 2 |  |
| --- | --- | --- | --- | --- |
|  | Total % of FN | % of FN by SNP Effect | Total % of FN | % of FN by SNP Effect |
| 50 | 33.9% | 83.8% | 21.2% | 99.5% |
| 100 | 11.2% | 99.1% | 3.2% | 100.0% |
| 200 | 0.9% | 100.0% | 0.1% | 100.0% |
| 300 | 0.0% | 0.0% | 0.0% | 0.0% |
| 600 | 0.0% | 0.0% | 0.0% | 0.0% |
| 700 | 0.0% | 0.0% | 0.0% | 0.0% |
| 800 | 0.0% | 0.0% | 0.0% | 0.0% |

FN = false negative. False negative rates were investigated by determining the number of significant and non-significant trials when comparing change in UPDRS over a simulation of two years after adding a "drug" effect that reduced UPDRS by 0.5 points per year. Total % of false negatives is the percentage of total iterations that resulted in a false negative. % of false negatives by SNP effect is the percentage of false negatives that were caused by the interaction effect on UPDRS progression by the rs9298897 and rs17710829 SNPs alone. Significance was determined by Z-score cut-off values of $\pm 1.96$ ( $p < 0.05$ ).

**Table 4.** False Positive Rates for Unbalanced Trials Over a Simulation of 2 Years

| Sample Size | Year 1 |  | Year 2 |  |
| --- | --- | --- | --- | --- |
|  | Total % of FP | % of FP by SNP Effect | Total % of FP | % of FP by SNP Effect |
| 50 | 6.1% | 49.2% | 5.6% | 71.4% |
| 100 | 5.8% | 82.8% | 5.2% | 84.6% |
| 200 | 5.9% | 72.9% | 5.7% | 86.0% |
| 300 | 4.5% | 71.1% | 5.7% | 86.0% |
| 600 | 4.5% | 71.1% | 5.7% | 75.4% |
| 700 | 6.0% | 78.3% | 4.8% | 91.7% |
| 800 | 5.7% | 75.4% | 4.3% | 74.4% |

FP = false positive. False positive rates were investigated by determining the number of significant and non-significant trials when comparing change in UPDRS over a simulation of two years. Total % of false positives is the percentage of total iterations that resulted in a false positive. % of false positives by SNP effect is the percentage of false positives that were caused by the interaction effect on UPDRS progression by the rs9298897 and rs17710829 SNPs alone. Significance was determined by Z-score cut-off values of $\pm 1.96$ ( $p < 0.05$ ).

### The Effects of Balancing on Variance per Simulated Trial Sample Size

We investigated the effects of balancing on GRS variance by applying an algorithm that distributed patients between the treatment and placebo arms in a way that minimized difference in genotype dosage between arms. The purpose of this was to explore a possible solution to large differences in GRS between randomly assigned arms, and to determine the scale of variance reduction after balancing patients. The pattern of decrease in GRS percent differences and variance with increasing sample size is still seen, however the magnitude of the differences across arms dropped substantially for all sizes as compared to the randomly assigned arms. Percent difference in mean GRS between arms at a sample size of 200 decreases significantly from 130.87% in significant iterations to 36.71%, a drop in GRS difference between groups of 94.16% (p = 9.7e-22, 95% CI [84.01, 104.31]) (Table 5).

**Table 5.** Average % Difference in Mean Balanced GRS

| Sample Size | Average % Difference in GRS (CI 95%) | % Difference Range (Min - Max) |
| --- | --- | --- |
| 200 | 36.71 (34.77 - 38.65) | 0.232 - 171.72 |
| 400 | 21.16 (20.12 - 22.20) | 0.003 - 199.09 |
| 600 | 14.88 (14.17 - 15.59) | 0.005 - 69.84 |
| 1,000 | 10.62 (10.11 - 11.13) | 0.026 - 58.90 |
| 2,000 | 6.11 (5.81 - 6.41) | 0.007 - 29.45 |
| 3,000 | 4.59 (4.37 - 4.81) | 0.004 - 27.42 |
| 4,000 | 3.40 (3.23 - 3.56) | 0.007 - 16.54 |
| 5,000 | 2.74 (2.60 - 2.87) | 0.002 - 16.65 |

The average percent difference between trial groups and related confidence intervals are shown for each sample size after the balancing algorithm had been applied. Minimum and maximum percent differences found between groups are also shown. These values are calculated using all 1000 iterations from each sample size.

While this is just one possible method to harmonizing trial arms, it is evidence that matching trial arms on genetic risk factors can allow for smaller sample sizes to be utilized in trials with less risk of false negatives. Component scores from the first three components of PCA were plotted to show the effect of balancing on the distribution of the scores across the components. The balanced trials are consolidated while the unbalanced trials are dispersed throughout the space, showing high variability in allele dosage. (Figure 3). To further investigate differences between balanced and unbalanced cohorts, the component scores from a PCA built using the variant dosage status of the 47 variants for each patient were used as predictors in logistic models to determine classification in the treatment or placebo arm. Nagelkerke’s pseudo R-squared was used to approximate variance explained by the model. R-squared for the unbalanced cohort was 18.8%, while R-squared for the balanced was lower at 12.4%. Average percent difference in mean GRS and variance in GRS was calculated using all 1000 iterations.

**Figure 3.**
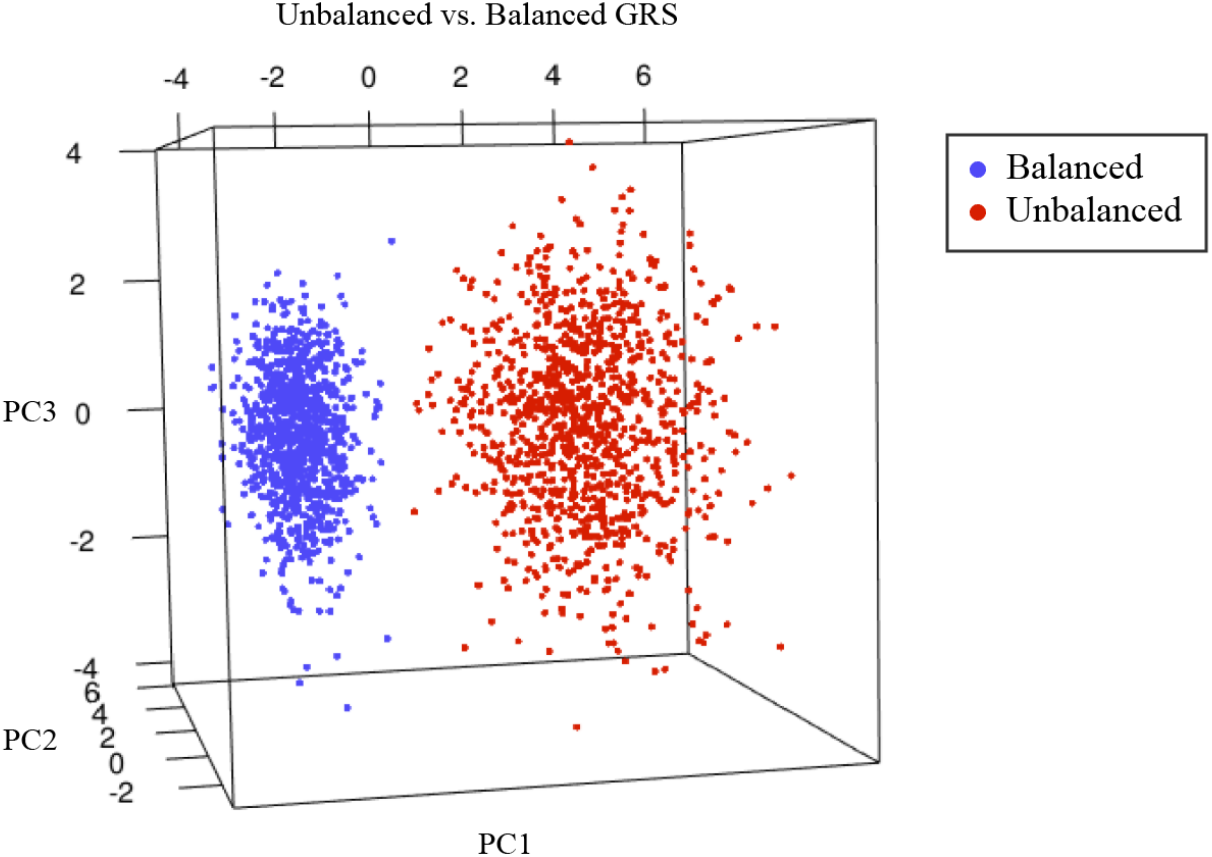
PCA component scores of balanced and unbalanced GRS for the main cohort. PCA was performed on unbalanced and balanced cohorts. Component scores from this analysis were used to visualize the effects of balancing on GRS at a sample size of 200.

### Genetic Heterogeneity in Randomized Virtual GBA Cohort and Balancing Effects

To investigate GRS variance within a stratified genetic subpopulation, we created a virtual cohort of *GBA* variant carriers using effect estimates and European population frequencies. We created an idealistic scenario in which all other variants are perfectly controlled, in order to demonstrate that randomization creates differences between patients and trial arms even with variants within a single gene. Variance analysis of the virtual *GBA* cohort revealed patterns similar to the larger cohort. Direct comparisons of the differences between the virtual and non-virtual cohort cannot be made, as the main cohort was ranked with the controls by Z-scores, while the virtual cohort lacked its own controls with which to create the same scale. However, direct comparison was not the goal, which was instead to investigate the pattern of variance in a subgroup of PD patients that are being enrolled in clinical trials. Analysis of differences in mean GRS between arms showed the same higher quantities of variance at small sample sizes. This virtual model only considers 3 known *GBA* variants with estimates of effect size. Real *GBA* cohorts that possess a wider range of variants both within and outside the gene are likely to display greater differences between trial arms. This demonstrates that even if other variants were perfectly controlled and patients were chosen by possessing a mutation in *GBA*, different *GBA* mutations are associated with different risk and effect, and this has the potential to cause differences between *GBA* trial arms. (Table 6) (Figure 2.B.). Balancing the *GBA* cohort produced results similar to those of the larger cohort as well, with the average percent difference in mean GRS between arms significantly decreasing from over 18.86% to 3.38%, a difference of roughly 15% (p = 1.5e-45, 95% CI [14.61, 16.35]). (Table 7).

**Table 6.** Average % Difference in Mean *GBA* GRS

| Sample Size | Average % Difference in GRS (CI 95%) Significant Iterations | Average % Difference in GRS (CI 95%) All Iterations | % Difference Range (Min - Max) |
| --- | --- | --- | --- |
| 50 | 18.86 (18.01 - 19.71) | 6.76 (6.43 - 7.08) | 0.000 - 28.78 |
| 100 | 13.31 (12.74 - 13.89) | 4.83 (4.61 - 5.05) | 0.000 - 20.78 |
| 200 | 9.63 (9.18 - 10.08) | 3.24 (3.09 - 3.39) | 0.000 - 14.41 |
| 300 | 7.59 (7.38 - 7.80) | 2.70 (2.57 - 2.83) | 0.013 - 9.50 |
| 600 | 5.62 (5.40 - 5.83) | 1.90 (1.81 - 1.99) | 0.001 - 8.12 |
| 700 | 5.31 (5.04 - 5.58) | 1.76 (1.67 - 1.84) | 0.001 - 7.67 |
| 800 | 4.77 (4.59 - 4.95) | 1.65 (1.58 - 1.73) | 0.006 - 6.43 |
| 1,000 | 4.49 (4.27 - 4.70) | 1.48 (1.41 - 1.55) | 0.003 - 7.77 |
| 2,000 | 3.09 (2.96 - 3.23) | 1.06 (1.01 - 1.11) | 0.001 - 4.42 |

The average percent difference between *GBA* trial groups and related confidence intervals are shown for each sample size for both significant iterations and total iterations. Significant iterations were determined by Z score cut-off scores of $\pm 1.96$ ( $p < 0.05$ ). Minimum and maximum percent differences found between groups for all 1000 iterations are also shown.

**Table 7.**
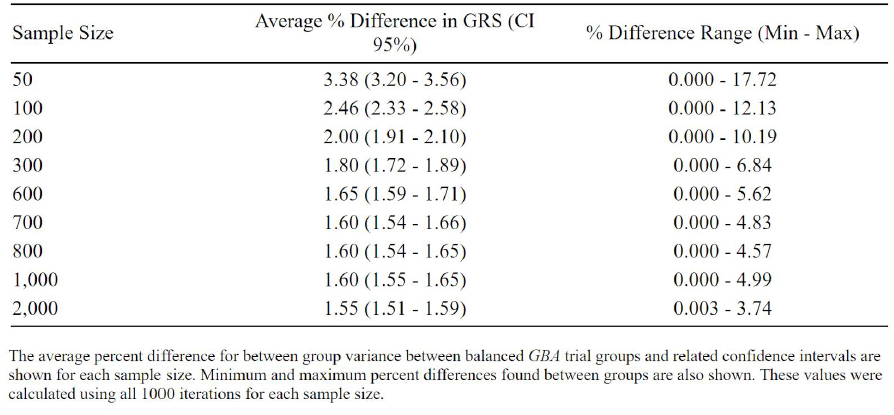
Average % Difference in Mean *GBA* Balanced GRS

### Genetic Heterogeneity in Terms of Heritability Estimates

We demonstrated the utility of balancing trials based on known genetic information by using PD heritability estimates as proxies. The “ceiling” values represent the high end of the estimated genetic variance that can be balanced in clinical trials, given the assumption that current genetic knowledge of PD explains a significant portion of disease variability. The “floor” values represent the low end of phenotypic variance estimates that can be balanced in clinical trials, given that the current heritability estimate of PD from GWAS is ∼26.9%. By triaging our estimates of cumulative genetic risk across groups by heritability (a proxy for penetrance of effect), we see similar trends but effects are slightly muted at small sample sizes. (Table 8).

**Table 8.** Average % Difference in Disease Readout, “Ceiling vs. Floor”

| Sample Size | "Ceiling" Average % Difference in GRS (CI 95%) | "Floor" Average % Difference in GRS (CI 95%) |
| --- | --- | --- |
| 200 | 130.87 (120.89 - 140.85) | 35.20 (32.52 - 37.89) |
| 400 | 120.59 (111.97 - 129.21) | 32.44 (30.12 - 34.76) |
| 600 | 91.31 (85.58 - 97.05) | 23.56 (23.02 - 26.12) |
| 1,000 | 71.15 (66.91 - 75.39) | 19.14 (18.00 - 20.28) |
| 2,000 | 48.03 (45.66 - 50.40) | 12.92 (12.28 - 13.56) |
| 3,000 | 39.00 (37.45 - 40.56) | 10.49 (10.07 - 10.91) |
| 4,000 | 35.02 (33.20 - 36.84) | 9.42 (8.93 - 9.91) |
| 5,000 | 29.87 (28.43 - 31.31) | 8.04 (7.65 - 8.42) |

The estimated range of percent difference in within group variance for each sample size is shown for significant iterations. Significant iterations were determined by Z score cut-off scores of $\pm 1.96$ ( $p < 0.05$ ). The minimum estimated values or "floor" values are calculated using current heritability estimates. The maximum estimate values or "ceiling" values assume a 1:1 relationship with variability in variants associated with Parkinson's and variance in disease readout.

## Discussion

Our simulation demonstrates that randomization without efforts to improve genetic imbalance, will result in significant variance between treatment and placebo arms in the vast majority of trials, of either a single SNP, multiple SNPs or GRS. Analysis of differences in mean GRS between arms in randomized simulated trials revealed that complete randomization of patients to each trial arm can result in large differences in variance between arms. As differences in GRS and allele carrier status can lead to differences in phenotypic readout, controlling this source of variance would lead to better balanced patients that could improve clinical trial results, depending on the genetic contribution to phenotype presentation. Virtual cohort analysis revealed that even when performing gene-specific stratification, different variants within a single gene can create large sources of variance between arms. Genetic balancing should be performed even in trials using small subgroups of variant carriers, such as *GBA* variant carriers, to mitigate the varying effects that different variants within and without the targeted gene can have on disease presentation, and thus, trial outcome. GRS has been found to be significantly associated with progression time to Hoehn and Yahr stage 3, with one standard deviation increase in total polygenic modeling GRS resulting in a 1.29 increase in hazard ratio.^30^ This finding, along with the results from Latourelle et al^10^, suggest that it is likely there are many more unknown associations between variants/GRS and progression in Parkinson’s phenotypes. While many of these effects are not yet known, it is probable that balancing both common and rare variants between trial arms can help to prevent possible effects on progression rates that may result in a higher variance in measurable outcomes like UPDRS score between patients by the end of the trial.

Single variant analysis showed that patient randomization tactics can cause significantly different variant distribution between trial arms. Given 47 variants with the possibility to affect phenotypic readout, there is only a roughly 10% chance that any given trial will have no differences in variant distributions between placebo and treatment arms. We found that while the number of significant variants fluctuated, there is an upward trend between number of significant variants and increasing sample size. This is most likely due to larger sample sizes having a greater chance of including relatively rare variants, such as the variants *LRRK2* p.G2019S and *GBA* p.N370S. Due to low population frequency (p.G2019: 0.0006% in Europeans; p.N370S: 0.0034% in Europeans)^22^, these variants are less likely to be picked up in smaller sample sizes, unless enriching for them. The low frequency of significance of these rare variants at small sample sizes is indicative of the high percent difference in GRS results being driven primarily by the more common variants. As sample size increases from 200 to 1000, the rarer variants have higher chances of creating significant differences in allele distribution between arms. While one or two patients with rare variants will not affect trial outcomes, an imbalanced group of large-effect rare variant carriers could skew results, particularly as misdiagnosis rate for neurodegenerative diseases is high. This is important to consider when designing a clinical trial, as larger sample sizes will decrease GRS variance, but may increase the chance of creating significantly different allele distributions. On the other hand, the magnitude of the percent difference in allele dosage decreases with increasing sample size, and as is seen in the false negative analyses, larger sample sizes will likely mitigate a rare variant imbalance unless the effect is very large. Genetic balancing would however, remove the need for making these decisions in regards to sample size, as it allows small samples to be utilized while removing a large source of variance.

Simulation analysis of a variant interaction effect on MDS-UPDRS showed how clinical trial outcomes could be affected by unbalanced variants with influence on phenotype progression. For small sample sizes, these effects are especially noticeable. As an example, considering the aforementioned 413 failed Alzheimer’s trials that took place between 2002 to 2012 and the simulated 2 year failure rate of 21.2% from the SNP interaction effect at n = 50, if the smaller trials had contained disparities in measurable disease outcome similar to the effects of rs9298897 and rs17710829 in Parkinson’s, roughly 87 of those trials may have failed due to genetic disease disparity. In addition, the interaction of the variants used in this simulation is quite rare, thus a similar effect size for more common SNPs and interactions will likely lead to higher rates of both false positive and false negative iterations. As this analysis was only based on the interaction between two SNPs, these results will vary in real clinical trials that hold more variability in terms of disease progression and genetic influences on phenotype. Whether false positive and false negative rates will be higher or lower will depend on the degree of imbalance and the effect size of the variant or interaction in question. Trial balancing is a fairly inexpensive tool that can be employed to prevent situations in which a large imbalance would affect results, as is seen in the interaction effect simulations.

While we have mainly discussed the effects of genetic variance in terms of clinical trial failure, differences in GRS between arms can also create a positive or negative bias for a drug. An important goal in a clinical trial is to determine if any witnessed benefits are a true drug effect, but variance in underlying genetics may cause false conclusions to be made. As an example of this effect, let us consider the following scenario. PD patients are randomly assigned to a placebo or treatment arm and MDS-UPDRS is chosen as the clinical readout to measure success of the trial. The trial is built based upon common trial inclusion criteria, such as base UPDRS and Hoehn and Yahr score. However, the control arm possesses alleles associated with said selection criteria, for which the alleles in question increase MDS-UPDRS progression by an additional 2 points per year. In this scenario, the drug is ineffective. However, the higher number of progression-affecting alleles in the control arm creates an observed lower MDS-UPDRS score in the treatment arm. This effect could be construed as the effect of the drug, when in fact, this is an example of “collider bias” or “selection bias”.^31^ This effect can occur in the opposite direction as well, such as in the case of an imbalance in carriers of the *LRRK2* G2019S, a mutation which has recently been connected to a slower rate of decline in motor functioning than in those without the mutation.^32^ Slower progression rates in the treatment arm could lead to false positives when the drug is ineffective. Balancing trials by GRS and allele distribution would control possible genetic bias that could lead to a false effect being classified as the effect of the tested drug.

Another source of variance that was not addressed in the current analysis, is genetic factors that may affect the metabolism of the drug itself. These will add to the overall genetic variance, and are harder to take into account when testing a new drug. In case of such variance, post-trial analysis of the treatment group can identify such variants, and a statistical correction could be applied based on the effects of such variants. A major limitation of such approach will be that a relatively large studies, or meta-analysis of studies, will be required.

It is important to note that power calculations for phase III clinical trials are based on phenotypic variance. As is demonstrated by the trials balanced on GRS, genetic matching in trials has the potential to reduce bias in randomized arms situationally, decreasing heterogeneity between arms to a degree that would theoretically need much larger sample sizes to accomplish. Reducing this heterogeneity allows for a reduction in sample size, cutting trial costs. This has been shown to be true in studies such as Stone et al.^33^, where they found that genotyping for *APOE* in AD trials resulted in an increase of power if sample sizes remained the same, or allowed for the number of study sites to be reduced without decreasing power.

While we have demonstrated the importance of genetic balancing across trial arms, an important caveat to consider is that we can only control variance to the extent of what is known by current heritability estimates. In addition to this study demonstrating the importance of genetic balance in clinical trials, it is also a call for larger scale genetic studies on progression that will allow us to account for and balance currently unknown sources of variance. Valuable future work would be to investigate the effects of variants on phenotypic outcomes that are measured by clinical trials, such as UPDRS, to gain further understanding of sources of variance in trial outcomes that can be controlled.

## Contributors

MAN and ZG-O conceived the study. MAN, ZG-O, and HL contributed to study design. MAN and HL contributed to data analysis. MAN, ZG-O, and HL contributed to data interpretation. MAN and CB contributed to data management and storage. HL drafted the manuscript and MAN, ZG-O, CB, LK, FF, HI, GF, AGD-W, DJS, IPDGC, and ABS performed critical review and additional writing. All authors gave approval for publication.

## Acknowledgments

See supplemental materials for IPDGC member acknowledgements.

